# RT-qPCR as a screening platform for mutational and small molecule impacts on structural stability of RNA tertiary structures

**DOI:** 10.1101/2022.01.16.476525

**Authors:** Martina Zafferani, Dhanasheel Muralidharan, Amanda E. Hargrove

## Abstract

The exponential increase in the discovery and characterization of RNA tertiary structures has highlighted their active role in a variety of human disease, yet often their interactome and specific function remain unknown. Small molecules offer opportunities to both decode these cellular roles and develop therapeutics, yet there are few examples of small molecules that target biologically relevant RNA tertiary structures. While RNA triple helices are a particular attractive target, discovery of triple helix modulators has been hindered by the lack of correlation between small molecule affinity and effect on structural modulation, thereby limiting the utility of affinity-based screening as a primary filtering method. To address this challenge, we developed a high-throughput RT-qPCR screening platform that reports on the effect of mutations and additives, such as small molecules, on the structuredness of triple helices. Using the 3’-end of the oncogenic non-coding RNA MALAT1 as an example, we demonstrated the applicability of both a two-step and a one-pot method to assess the impact of mutations and small molecules on the stability of the triple helix. Employment of a functional high-throughput assay as a primary screen will significantly expedite the discovery of probes that modulate RNA triple helices structural landscape and, consequently, help gain insight into the roles of these pervasive structures.

## Introduction

Advances in biophysical techniques aimed at studying RNA structure have resulted in an exponential increase in discovery and characterization of RNA tertiary structures, including RNA triple helices.^1, 2^ The recent surge in interest and characterization of RNA triple helices across various kingdoms of life has led to coining of the term ‘triplexome’, referring to the diverse and large number of RNA triple helices involved in cellular processes.^3^ While many of the functions of triplexome members are still being elucidated, the recognized roles of this RNA structural topology include protection of RNA from degradation, protein recruitment, and nuclear localization. RNA triple helices have been found in a variety of long non-coding RNAs, and the structuredness of the triplex motif has been shown to render the transcript refractory to exonuclease degradation, ultimately promoting its cellular accumulation.^3–5,4^ Despite the continuous increase in the size of the triplexome, small molecule targeting of these structures has significantly lagged, with only a few examples published.^5–7^ Notably, recent studies showed that small molecule modulation of RNA structure and function cannot always be predicted by affinity alone, highlighting the need for cost efficient high-throughput assays that report on the effects of small molecule:RNA interactions in relationship to structural stability.^8^ Furthermore, affinity-based screening platforms for RNA tertiary structures are limited due to the inherent challenges of optimizing binding assays for larger, more complex structures.^8^ Function-based assays would also help bridge the gap between *in vitro* screening and biological activity, thereby expediting the discovery of small molecule modulators for the continuously growing number of RNA triple helices. In turn, affordable high-throughput screening platforms can enable researchers to screen both mutations and small molecules and assess their impact on RNA structural stability, ultimately providing essential insight into the role of complex RNA structures in diseased pathways.

Methodologies commonly employed to assess RNA structural stability include circular dichroism (CD), UV absorbance spectroscopy (UV-Vis), differential scanning calorimetry (DSC), and differential scanning fluorimetry (DSF).^9–12^ While CD and UV-Vis can provide a variety of thermodynamic parameters, they suffer from low throughput and generally require large amounts of RNA, severely limiting their scope and application to sizeable screenings. DSC and DSF can be optimized for high-throughput screening and can provide information on the impact of a variety environmental factors on RNA structure, but the use of heat can lead to results that do not reflect an output relevant to biological environments.^12^ Furthermore, while DSC and DSF can readily identify molecules with thermal stabilizing effects, the identification of molecules with destabilizing effects is rare. Finally, recent developments and optimization of RNA enzymatic degradation assays have shown them to be uniquely poised to interrogate the effect of additives on RNA structural stability and resistance to enzymatic degradation in a more biologically relevant setting.^5, 6^ However, enzymatic degradation assays are often analyzed through gel electrophoresis and are thus severely limited both in throughput and quantitative potential. It is imperative to develop high-throughput methodologies that assess the effect of small molecules and other additives on RNA structure reliably, quickly, and cost-effectively under biologically relevant conditions.

While the link between structural stability of RNA triplexes and their biological function is a relatively recent discovery, the stability-function relationship is a well-established relationship for RNA G-quadruplexes (rG4s), making them a good reference for our functional assay.^13^ For example, highly stable rG4s can result in a premature stop in ribosomal scanning and prevent translation of the mRNA transcript downstream.^13^ This finding led to the preliminary investigation of quantitative PCR (qPCR) on the reverse transcribed cDNA template as a potential method to identify G-quadruplex structures.^14, 15^ Recently, Katsuda and co-workers were able to employ this method as a secondary screen to identify small molecules that increased the stability of RNA G-quadruplexes by observing their effect on the length and amount of cDNA formed during reverse transcription of the RNA.^16^ The small molecule identified as an inhibitor of the elongation reaction of the reverse transcriptase (RT) in the mRNA TERRA was also confirmed as translation inhibitor *in cellulo*, corroborating the biological relevance of the *in vitro* RT assay.^16^ Based on the success of this method in identifying functional modulators of a G-quadraplex, we sought to ask whether an RT-qPCR reaction could be optimized for application in a general cost-effective high-throughput platform for RNA tertiary structural stability. In addition to the use of a two-step method and high input of RT in the previous experiments, the reported structuredness and thermal stability of RNA G-quadruplexes brings into question whether this assay can be optimized and applied to more complex and/or less stable RNA tertiary structures.^16^ We also sought to assess whether an RT-qPCR method could identify both stabilizing and destabilizing mutations and small molecules. We hypothesized that such an assay could both elucidate key interactions necessary for triplex formation and stability via mutant analysis and provide useful modulators to facilitate elucidation of triplex-mediated regulatory pathways and unravel new therapeutic avenues.

To ensure the applicability and biological relevance of the newly developed high-throughput RT-qPCR screening platform presented herein, we chose the well-studied MALAT1 triple helix as our case-study. The A-U rich blunt-ended MALAT1 triple helix forms at the 3’-end of a 6.7 kb long non-coding RNA (lncRNA) found to be overexpressed in several types of cancers and reported to be implicated in a plethora of human diseases, including diabetes.^17, 18^ While the structure and functions of the entire transcript are still under investigation, the triple helix has been reported as essential for protection of the transcript from enzymatic degradation, ultimately leading to increased cellular accumulation of MALAT1.^18, 19^ Indeed, knockdown of MALAT1 in small cell lung adenocarcinoma mouse models led to significant decrease in tumor size and metastasis, confirming the oncogenic role of the lncRNA MALAT1.^20^ Biophysical studies by Steitz and coworkers further showed that mutations aimed at destabilizing the triple helix structure led to a significant depletion of MALAT1 *in cellulo*, establishing the modulation of the MALAT1 triplex as a potential therapeutic avenue.^18^

## Results and Discussion

### Assay and MALAT1 construct design

The 3’-MALAT1 triple helix forms via recruitment of a genomically encoded A-rich tail to the adjacent U-rich region after processing of the full-length transcript.^21^ Given the critical function and sequestration of the 3’-end, we sought to avoid the use of an A-tail specific reverse primer for the reverse transcription reaction. We thus synthesized a construct containing a primer handle commonly used in chemical probing experiments (SHAPE cassette, **Figure 1**, orange) that has been reported to not affect structures of RNA transcripts under investigation.^22, 23^ Indeed, SHAPE cassettes have been successfully employed in chemical probing experiments of MALAT1 and MALAT1-like evolutionarily conserved transcripts, and the results were consistent with triple helix formation (**Figure S1**).^24^ In the system designed here, small molecules that stabilize the triple helix will result in inhibition of the reverse transcription reaction, due to the inability of the chosen RT enzyme (SSIV, ThermoFisher) to unwind structured regions. Inhibition of the elongation reaction will ultimately result in lower levels of cDNA produced, which can be measured quantitatively via qPCR (**Figure 1**). Analogously, small molecules that destabilize the triple helix conformation will enable for more efficient readthrough of the RT, yielding higher cDNA and a resulting lower C_t_ value than the control (**Figure 1**).

**Figure 1.**
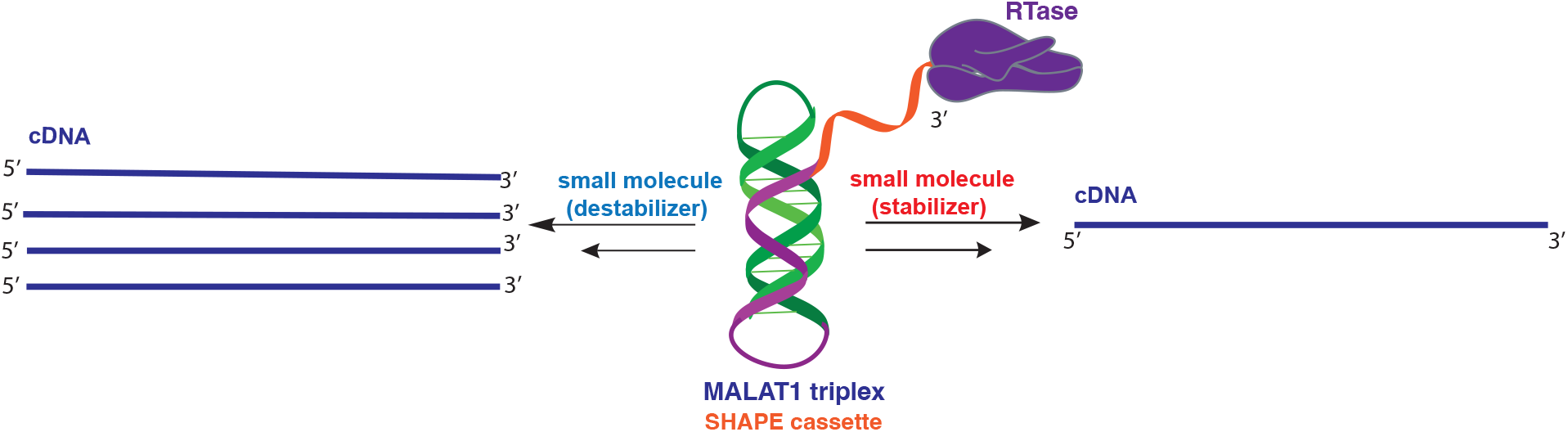
Schematic representation of small molecule-induced structural modulation of the MALAT1 triple helix assessed via RT-qPCR assay. The MALAT1 triple helix (green/purple) is equipped with a structural SHAPE cassette (orange) to prevent competition between primer binding and triplex formation. According to the assay design, small molecules that destabilize the triple helix construct result in lower frequency of RT stalling and, consequently, in more full-length cDNA synthesis (*left*). Small molecules that stabilize the triple helix structure result in higher occurrence of RT stalling, ultimately resulting in lower amounts of full cDNA synthesis (*right*). Reverse transcription reactions are then followed by qPCR for quantification (not shown).

Upon successful synthesis of the designed MALAT1 construct (**Table S1, Figure S1**), we first optimized the assay for the evaluation of mutational effects on structural stability, particularly destabilizing effects. Accordingly, we synthesized a construct containing a MALAT1 mutant where one of the uracil involved in a base triple within the triplex core is mutated to a cytosine (U13C). This U13C mutant was reported by Steitz and co-workers as a triplex destabilizing mutation that resulted in a decrease in full-length MALAT1 levels *in cellulo* (**Figure 2**).^18^

**Figure 2.**
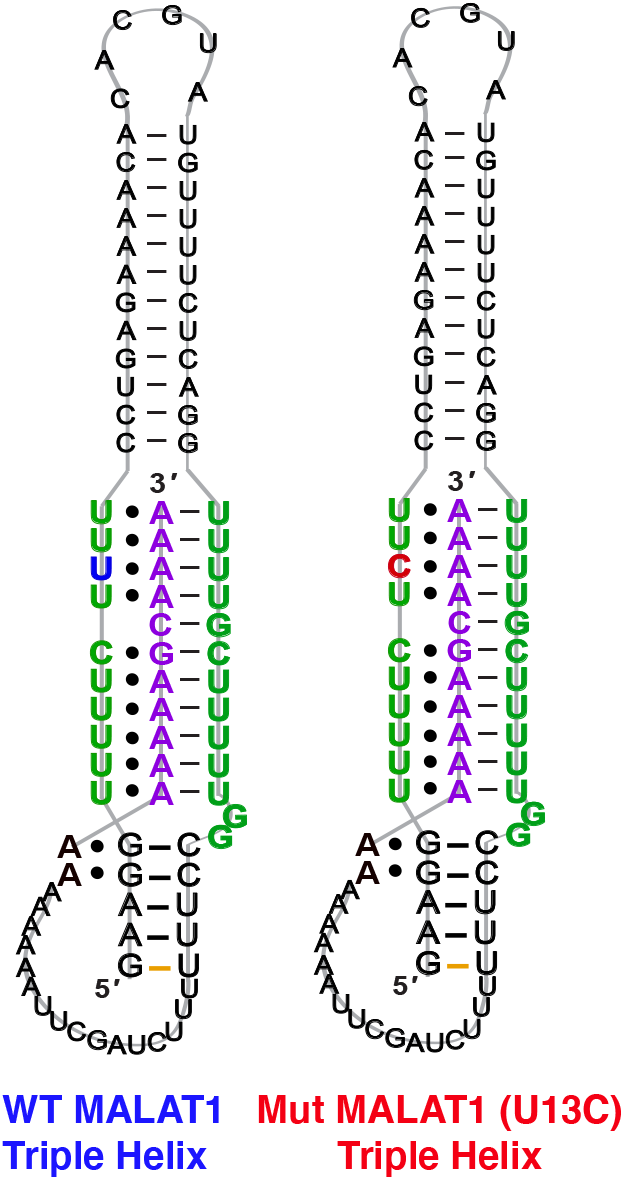
2D structure and sequence of MALAT1 WT and U13C mutant.

### Two and one-step RT-qPCR

A two-step RT-qPCR was first utilized to allow for optimization of the reverse transcription and qPCR steps independently. The MALAT1 wild type (WT) and mutant (U13C) were incubated with DMSO control at room temperature after which reverse transcriptase (SSIV), magnesium, primers, and dNTPs were added on ice. The reverse transcription reaction was then carried out at 37°C and inactivated by heating to 98°C after 15 minutes from the start of the reaction. The reaction was then aliquoted in a 96-well light cycler plate and qPCR mix was added to each well and placed in a real-time qPCR machine for cDNA amplification (**Figure 3A**). Both reverse transcription and qPCR steps were optimized to obtain WT amplification with a C_t_ that would allow detection of ΔΔC_t_= ± 4 without exceeding the instrument detection limit (< 2 C_t_) or incurring non-specific amplification (> 25 C_t_).^25^ Detection of ΔC_t_= ± 2 corresponds to a 75% change in RT efficiency.^16^ Initial RNA quantities were in line with manufacturer recommendations. The reverse transcription reaction was optimized by testing whether the reaction needed the presence of “first-strand buffer” for reverse transcription (1M DTT, 0.5M Tris-HCl, 0.25 M KCl, at pH 8.0). Two reverse transcription reaction were carried out in small scale (20μL) with first strand buffer or water and the reaction was quenched and cleaned up after 30 minutes at 37°C. The presence of cDNA was assessed by Agarose gel and by Nanodrop, which revealed equal amounts of cDNA in both reactions. Given the presence of denaturant and the non-biologically relevant pH of the first strand buffer, the RT reaction was subsequently optimized without first strand buffer. Subsequent optimization included variation of reverse transcriptase amount (Thermo Fisher, 30-150 units), magnesium chloride concentration (1-7.5 nM), reverse primer concentration (150-500 nM), dNTP concentration (100-600 nM), and RNA concentration (10-15nM). After successful optimization of the reverse transcription reaction (final conditions: 150 SSIV units, 10nM of RNA, 150nM of primers, 600 nM of dNTPs, 300 nM of MgCl_2_), qPCR (KAPA) was optimized by varying the amount of cDNA added to the mix (1-3μL of a max 200 ng/μL RT reaction). Analysis of the qPCR curve yielded a lower C_t_ value for MALAT1-U13C when compared to the MALAT1-WT, ultimately yielding a ΔC_t_= 2.1 ± 0.2 **(Figure 3B-C**).

**Figure 3.**
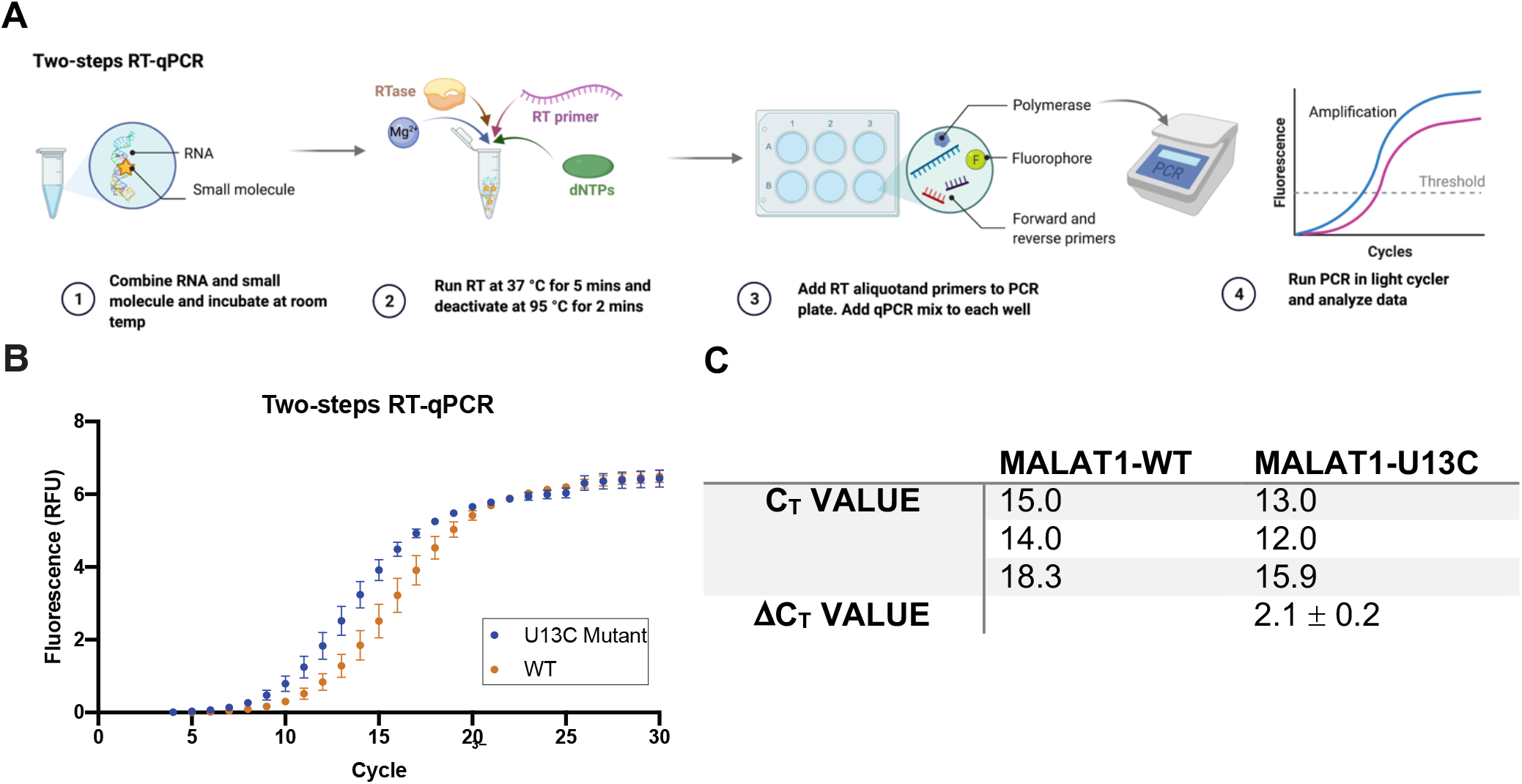
Two-step RT-qPCR system. (A) Schematic of the two-step RT-qPCR reaction with the reverse transcription reaction being performed in a thermocycler and then being aliquoted in a 96-well plate for qPCR amplification, which is performed in a light cycler. (B) Amplification curves obtained from 3 independent replicates of RT-qPCR of the U13C MALAT1 mutant and the WT triple helix. Error bars are standard deviation calculated over the three independent experiments. (C) C_t_ values calculated over three independent experiments for WT and U13C mutant (ΔC_t_ value =C _t Mut_ – C _t WT_). The mutated destabilized construct gets amplified faster than the WT, in agreement with trends reported by Steitz and co-workers.

Having optimized a two-step reaction, the MALAT1 WT and Mut were analogously employed in a one-pot RT-qPCR kit (QuantaBio), which would enable cost-efficient, high-throughput screening of additives and small molecules. In this system, both the reverse transcription and the qPCR amplification are performed directly in a real-time qPCR instrument (**Figure 4A**). The one-pot reaction was optimized by varying the RNA concentration in the reaction (1-20 nM) and the forward and reverse primer concentration (200-500 nM) in the same buffer previously used to obtain C_t_ values in the same range as the two-step RT-qPCR optimized above. Once again, we found that the MALAT1 mutant has lower C_t_ value than the wild type, yielding a ΔC_t_= 2.2± 0.6. In summary, both methodologies were confirmed to report on mutation-induced changes in MALAT1 triplex structural stability and, most importantly, the trends observed corroborate those which Steitz and co-workers observed in a cellular environment.

**Figure 4.**
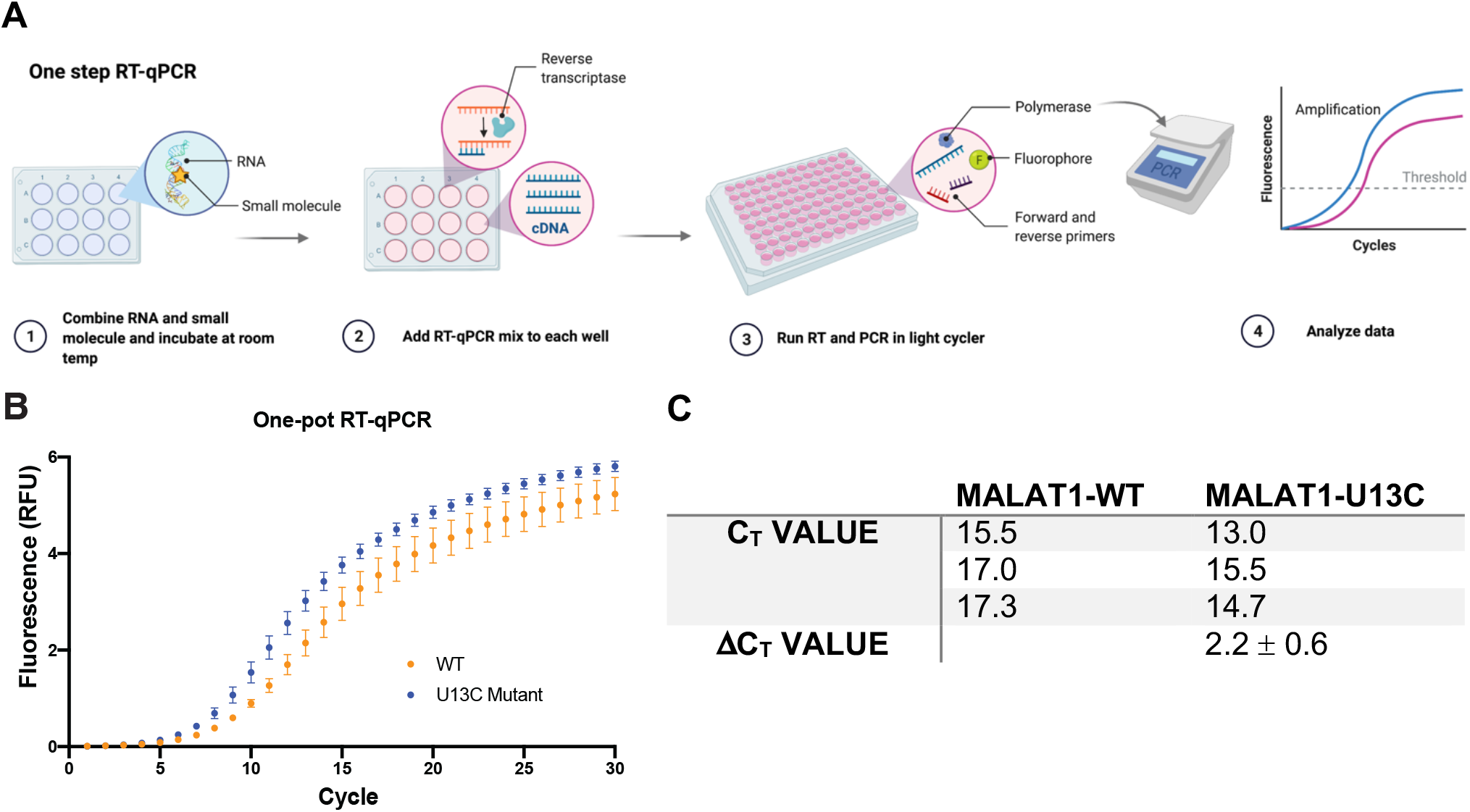
One pot RT-qPCR system. (A) Schematic of the one-pot RT-qPCR reaction with reverse transcription reaction being performed a 96-well format in a light cycler instrument. (B) Raw data obtained from 3 independent replicates of RT-qPCR of the U13C MALAT1 mutant and the WT triple helix. Error bars are standard deviation calculated over the three independent experiments. (C) C_t_ values obtained for the Mut and WT constructs (ΔC_t_ value =C _t Mut_ – C _t WT_). The mutated destabilized construct gets amplified faster than the WT, in agreement with trends reported by Steitz and co-workers.

### Small Molecule Screening

Next, we tested the applicability of both RT-qPCR routes to evaluate the impacts of small molecules on the MALAT1 WT triple helix. For this purpose, we chose two previously published small molecules that have been classified as stabilizers and destabilizers, respectively (**Figure 5A**).^5, 6^

**Figure 5.**
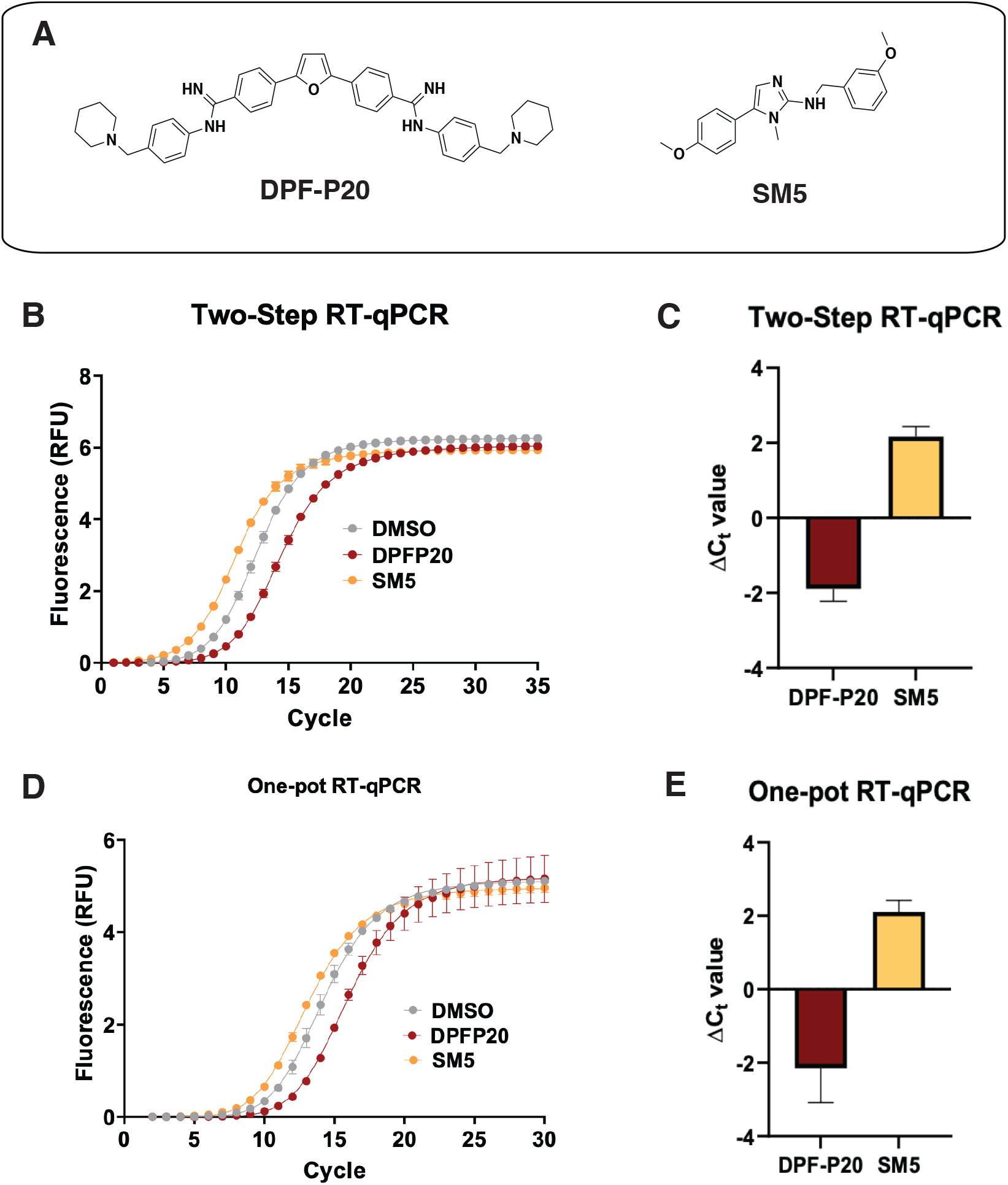
Application of RT-qPCR assays to assess small molecule effect on MALAT1 WT triple helix stability. (A) Structures of two small molecules chosen for assay validation. Both DPF-P20 (a MALAT1 triplex stabilizer) and SM5 (a MALAT1 triplex destabilizers) were previously evaluated in relationship to their effect on triplex enzymatic degradation.^5, 6^ (B) Raw data obtained for the two-step RT-qPCR procedure across for both small molecules. (C) Small molecule ΔC_t_ values calculated in reference to DMSO from 4 independent replicates in reference to DMSO are in agreement with their reported effect on MALAT1 triplex enzymatic degradation. (D) Raw data obtained for the one-pot RT-qPCR procedure for both small molecules. (E) Small molecule ΔC_t_ values calculated in reference to DMSO from 4 independent replicates in reference to DMSO are in agreement with their reported effect on MALAT1 triplex enzymatic degradation and in line with the values obtained in the two-step RT-qPCR procedure. All error bars represent the standard deviation calculated over the four independent experiments.

**DPF-P20** has been recently reported as a MALAT1 triple helix stabilizers as it increases the triplex thermal stability measured via DSF as well as inhibiting RNase R-mediated exonucleolytic degradation of the triplex (**Figure 5A**).^5^ Both the two-step and the one-pot method identified **DpF-P20** as a triplex stabilizer, yielding less cDNA and higher C_t_ values than the DMSO control (ΔC_t (2step)_ = −1.9 ± 0.3 ΔC_t (1pot)_ = −2.3 ± 0.7) (**Figure 5C-E**). Recently reported by Le Grice and coworkers, **SM5** was identified as a destabilizer of the MALAT1 triplex resulting in reduction of MALAT1 accumulation in *ex vivo* organoids breast cancer models (**Figure 5A**).^6^ **SM5** has also been shown by Donlic and co-workers to increase RNase R-mediated exonucleolytic degradation of the triple helix over time.^5^ In line with previous studies, **SM5** resulted in lower C_t_ values than the DMSO control, confirming its triplex destabilizing properties (ΔC_t (2step)_ = 2.2 ± 0.3 ΔC_t (1pot)_ = 2.1 ± 0.3) (**Figure 5B-C**). ΔC_t_ values were in the same range in both the two-step and the one-pot RT-qPCR method, showcasing the reliability and applicability of both approaches. (**Figure 5D-E**). As expected, given the sensitivity of RT-qPCR, Z-factor control experiments for the one-pot method were suitable for a high-throughput screening platform (*Z-*factor = 0.93, **Figure S3**). The |ΔC_t_| values observed for the MALAT1 triplex modulators are comparable to the values obtained by Katsuda and co-workers for the best leads of G-quadruplex stabilizers identified. The authors used ΔC_t_ ≥ 2 as a cut off for lead molecules since a value of 2 corresponded to a 75% decrease in RT elongation.^16^ Both small molecules, **DPF-P20** and **SM5**, would be classified as hits under these conditions, and we propose the same cut off for the assay reporter here.

## Conclusion

Here, we report the development and optimization of a new high-throughput screening platform that assesses the effects of mutations and small molecule additives on the structural stability of the MALAT1 3’-end RNA triple helix, one of the best studied disease-relevant RNA triple helices. While recent efforts aimed at small molecule targeting and modulation of this RNA motif have identified small molecule binders via screening, the effect of the reported small molecules on MALAT1 triplex stability have been studied through low throughput and/or non biologically relevant techniques.^5, 7^

In this work, we developed a high-throughput RT-qPCR screening platform, accessible via both two-step and one-pot protocols, utilizing the MALAT1 triple helix and a biologically relevant mutant construct. Robust differences in C_t_ values of the U13C mutant relative to wild type recapitulated the destabilizing effects of the point mutation, which was previously reported to lead to a decrease of MALAT1 transcript accumulation *in cellulo*. These findings underscored the applicability of this platform to evaluate the effects of mutations on structural stability as well as the likely biological relevance of the trends observed. We then chose two MALAT1 small molecule ligands previously published as stabilizers or destabilizers, respectively, and evaluated their impacts on the MALAT1 triple helix in both methods. Once again, both approaches resulted in trends consistent with previously published effects of the small molecules on RNase R-mediated exonucleolytic degradation of the triplex.

The RT-qPCR-based screening method developed herein establishes a high-throughput platform that can identify RNA-targeted small molecules that have both stabilizing and destabilizing effects on RNA tertiary structure. The ability to identify probes with opposite impacts can greatly help elucidate the many biological roles of RNA tertiary structures such as triple helices in human disease and expedite the discovery of RNA-targeted therapeutics. Having access to a cost-efficient high-throughput structural stability screening platform can significantly increase the ability to evaluate small molecule selectivity for one structure over another in relationship to structural stability. In turn, the data gathered can help address several unanswered questions such as what molecular properties make a small molecule a stabilizer or a destabilizer and whether we can use structural stability data as a guiding principle for small molecule design and synthesis. We expect that answering these standing questions will move the scientific community toward the efficient development of RNA-targeted small molecule therapeutics.

## Experimental Procedures

### Synthesis of RNA constructs

DNA template sequence was purchased from Dharmacon and forward and reverse primers were purchased from Integrated DNA technologies (IDT) (**Table S1**). For PCR amplification the following reagents were added to for a given 50μL final reaction volume. First, the entire working space was sprayed with RNase Zap to prevent any contamination. Next, in the desired amount of sterile PCR tubes (ThermoFisher) 12.5 μL of RNase free water and 12.5 μL of Q5 reaction buffer were added, followed by 1μL of dNTPs (10mM), 2.5 μL of forward and reverse primer (10μM), 1μL of DNA template (50ng/μL) and, lastly 0.5 μL of Q5 polymerase. The DNA template was then amplified for 30 cycles in an Eppendorf Echo thermocycler. A Zymo DNA-clean-up kit was then utilized to clean-up the desired DNA sequence. A solution of amplified DNA in water was made to reach 28-35 ng/μL. The sequence was then in vitro transcribed (IVT) using the following protocol. For a given 50μL IVT reaction the following were added in respective order: 31.75 μL of RNase free water, 1.25μL MgCl_2_ (1M), 2μL Tris-HCl (1M at pH 8.0), 1.25μL of spermidine (0.1M), 0.5 μL of Triton-X (0.1%), 0.5 μL of DTT (1M), 0.2 μL of pyrophosphatase enzyme (100U/mL), 5 μL of DNA template and, finally, 2.5 μL of T7 polymerase enzyme kindly provided by the Tolbert lab at Case Western University. The reaction is then incubated at 37°C for 12 hours. Following incubation, the reaction is treated with DNase I and DNase I buffer twice in intervals of 30 minutes, followed by addition of 10% of the reaction volume of EDTA. The desired RNA is then extracted using phenol chloroform extraction and further purified via ethanol precipitation. Purity and size of the RNA construct is confirmed by Small RNA chip on Agilent Bioanalyzer (**Figure S2**).

### Two Step RT-qPCR

RNA was synthesized and purified as previously described. For a given reverse transcription reaction 10 nM of RNA were incubated in water with DMSO or 10 μM of small molecules at room temperature for 20 mins. After incubation the PCR strip was placed on an ice block. A mastermix of reverse transcription reaction reagents was prepared and aliquoted in each reaction to reach a total of 20 μL yielding a final concentration of 150 SSIV (ThermoFisher) units, 150nM of primers, 600 nM of dNTPs (BioRad), 300 nM of MgCl_2_ for a single reaction. The PCR strip was then incubated at 37 °C in a thermocycler (Eppendorf, Nexus Gradient) for 15 minutes before inactivating the RT enzyme for 5 minutes at 98 °C. The PCR strip was then placed on an ice block. In the meantime, a qPCR mix was prepared by adding 0.4 μL of 10 μM of both forward and reverse primer solution, 10 μL of SYBR qPCR mix (KAPA SYBR fast qPCR kit), 18 μL of nuclease free water. A total of 24 μL of the qPCR mastermix was aliquotted in each well of a 96-well lightcycler plate (Roche 96) and 1 μL of RT reaction was aliquoted in each well. Amplification protocol was run by incubating at 95 °C for 3 minutes followed by a 3-step amplification for 25-45 cycles and a final cooling to 40 °C over the course of 10 minutes. Results were analyzed using the Roche 96 light cycler software v1.1.

### One Pot RT-qPCR

RNA was synthesized and purified as previously described. For a given RT-qPCR reaction 10 nM of RNA were incubated in water with DMSO or 10 μM of small molecules at room temperature for 20 mins in a 96 well lightcycler plate (Roche). During incubation an RT-qPCR mastermix was prepared by adding 25 μL of 1 step SYBR mastermix (QuantaBio), 1 μL of qScript RT enzyme (QuantaBio 1-step RTqPCR kit), 2.5 μL of a 10 μM forward and reverse primer solution. The mastermix of reverse transcription reaction reagents was prepared and aliquoted in each well by adding 31 μL of the master mix to each RNA-DMSO or RNA-Small molecule well. The plate was then sealed with optically clear foils and inserted in a Roche 96 light cycler. The RT step was performed by incubating at 48 °C for 10 minutes, followed by an inactivation at 95 °C for 5 minutes. The qPCR step immediately followed with a 3-step amplification carried for 30-35 cycles, which was followed by cooling to 40 °C over the course of 10 minutes. Results were analyzed using the Roche 96 light cycler software v1.1.

## Acknowledgements

We are grateful to current and past Hargrove lab members, especially to Dr. Emily McFadden and Dr. Sarah Wicks, for their feedback and input. We are grateful to the Tolbert lab for their generous donation of the T7 polymerase used for *in vitro* transcription. We would also like to thank Dr. Jessica Brown for kindly sharing illustrator files to make 2D-renderings of the MALAT1 triple helix and the Duke Biology Department for sharing the use of their Roche light cycler instrument.

