## Supplemental Information for "RT-qPCR as a screening platform for mutational and small molecule impacts on structural stability of RNA tertiary structures"

***Table S1**. Nucleic acid sequences for synthesis of the MALAT1 construct with the addition of the 3'-end SHAPE cassette (shown in red)*

| *Name* | Sequence |
| --- | --- |
| *MALAT1-WT DNA template* | GGAAGGTTTTTCTTTTCCTGAGAAAACAACACGTATTGTTTTCTCAGGTTTTGCTTTTTGGCCTTTTTCTAGCTTAAAAAAAAAAAAAGCAAAATCGATCCGGTTCGCCGGATCCAAATCGGGCTTCGGTCCGGTTC |
| *MALAT1-WT Forward Primer* | GAAATTAATACGACTCACTATAGGAAGGTTTTTCTTTTCCTGAGAAAACAACACGTATT |
| *MALAT1 SHAPE cassette Reverse Primer* | mG[mA]ACCGGACCGAAGCCC |
| *MALAT1-U13C DNA template* | GGAAGGTTTTTCT**C**TTCCTGAGAAAACAACACGTATTGTTTTCTCAGGTTTTGCTTTTTGGCCTTTTTCTAGCTTAAAAAAAAAAAAAGCAAAATCGATCCGGTTCGCCGGATCCAAATCGGGCTTCGGTCCGGTTC |
| *MALAT1-U13C Forward Primer* | GAAATTAATACGACTCACTATAGGAAGGTTTTTCTCTTCCTGAGAAAACAACACGTATT |

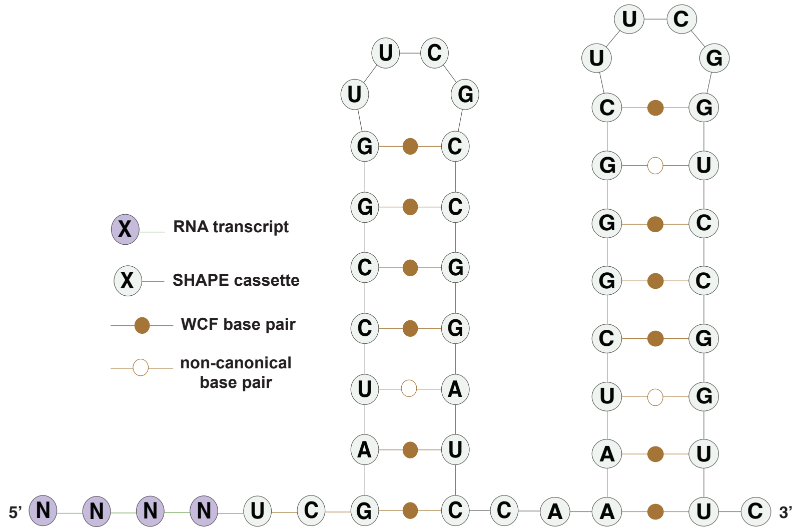


**Figure S1.** Representation of structure and sequence of the SHAPE cassette (orange) as reported by Wilkinson and co-workers.[1]

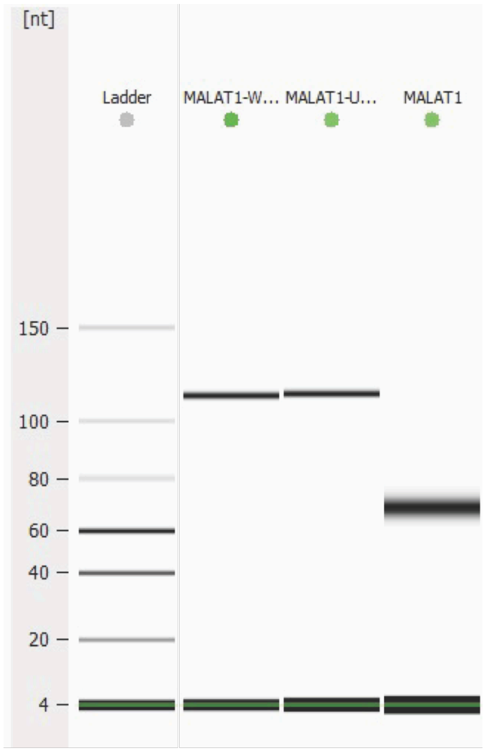


**Figure S2.** MALAT1-WT and MALAT1-U13C triple helix construct run on Small RNA chip on Agile bioanalyzer. Construct size is within 25% confidence value of sizing for the gel chip.

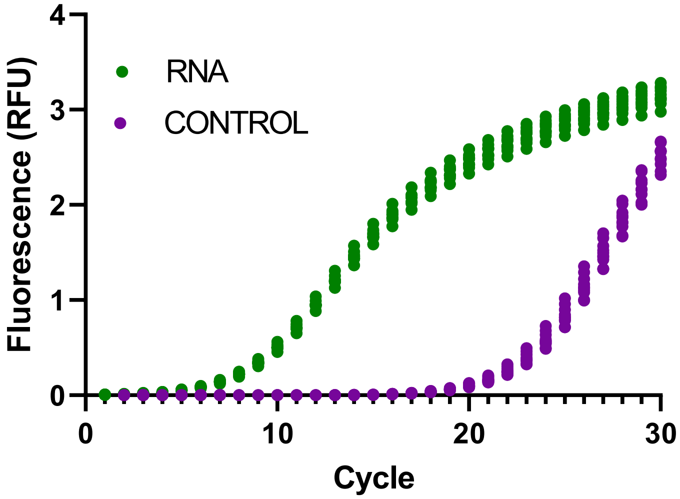


**Figure S3.** Raw curve of z'-factor experiment with amplification of MALAT1_WT or control. The experiment was run in two independent replicates with 10 wells of RNA and 10 wells of controls (no RNA) each. Z-factor was calculated as previously reported.[2]
